# Influence of isolation by environment and landscape heterogeneity on genetic structure of wild rice *Zizania latifolia* along a latitudinal gradient

**DOI:** 10.1101/2020.05.29.124685

**Authors:** Godfrey Kinyori Wagutu, Xiangrong Fan, Wenlong Fu, Wei Li, Yuanyuan Chen

**Affiliations:** Key Laboratory of Aquatic Botany and Watershed Ecology, Wuhan Botanical Garden, Chinese Academy of Sciences, Wuhan, China; University of Chinese Academy of Sciences, Beijing, China; Sino-Africa Joint Research Center, Chinese Academy of Sciences, Wuhan, China; College of Science, Tibet University, Lhasa, Tibet Autonomous Region, China; Research Center for Ecology and Environment of Qinghai-Tibetan Plateau, Tibet University, Lhasa, Tibet Autonomous Region, China

**Keywords:** genetic structure, isolation by environment, landscape heterogeneity, latitudinal gradient, microsatellite markers, *Zizania latifolia*

## Abstract

Global aquatic habitats are undergoing rapid degradation and fragmentation as a result of land-use change and climate change. Understanding the genetic variability and adaptive potential of aquatic plant species is thus important for conservation purposes. In this study, we investigated the role of environment, landscape heterogeneity and geographical distance in shaping the genetic structure of 28 natural populations of *Zizania latifolia* (Griseb.) Turcz. Ex Stapf in China based on 25 microsatellite markers. Genetic structure was investigated by analysis of molecular variance (AMOVA), estimation of *F*_ST_, Bayesian clustering and Thermodynamic Integration (TI) methods. Isolation by environment (IBE), isolation by resistance (IBR) and isolation by distance (IBD) hypotheses were compared using a reciprocal causal model (RCM). Further, generalized linear models and spatially explicit mixed models, by using geographic, landscape and genetic variables, were developed to elucidate the role of environment in driving *Z. latifolia* genetic diversity. The genetic differentiation across all populations was high: *F*_ST_ = 0.579; Ø_pt_ = 0.578. RCM exclusively supported IBE in shaping genetic structuring, only partial support for IBR, but not for IBD. Maximum temperature of the warmest month and precipitation seasonality were the plausible parameters responsible for genetic diversity. After controlling for spatial effect and landscape complexity, precipitation seasonality was significantly associated with genetic diversity. Based on these findings, genetic structure of *Z. latifolia* across China seem to be as a result of local adaptation. Environmental gradient and topographical barriers, rather than geographical isolation, influence genetic differentiation of aquatic species across China resulting in instances of local adaptation.

## 1 INTRODUCTION

Species adaptation to the changing environment mainly depend on the level of standing genetic variation in the populations (Lande and Shannon, 1996; Bell and Collins, 2008). Gene flow patterns, one of the factors influencing genetic variability, are affected by the movement of gametes, individuals and groups of individuals under the influence of landscape features (Sork et al, 1999). This further impacts the spatiotemporal patterns of genetic structure and evolution of natural populations (Slatkin, 1985). Climate and land-use changes are the major drivers of environmental alterations. For instance, global warming has led to temperature and precipitation fluctuations with instances of extreme episodes (Grant et al., 2017; Marrot et al., 2017), which result in plant populations’ selective pressure (Hoffmann and Sgrò, 2011; Cao et al., 2015). On the other hand, increase in human population has resulted in rapid conversion of natural lands to farmlands, urban and industrial areas leading to habitat loss and fragmentation (Zhao et al., 2013; Davidson, 2014). The overall effect is decline in population size of native species, which is the initial stage of species local disappearance and eventual extinction (An et al., 2007; Qin, 2008; Halley et al., 2016; Pykälä, 2019). Habitat degradation has profound effects particularly on aquatic plants that live in fragmented islands within the terrestrial landscapes. Additionally, most aquatic plants persist as meta-populations, the long-term survival of which depends on continuous gene flow among populations (Barrett et al., 1993; Santamaría, 2002). Therefore, clear understanding of genetic variability of aquatic plant species occurring across a large spatial scale is important for estimation of the current vigor under the different abiotic and biotic conditions and hence the potential to adapt to the changing environment (Catullo et al., 2019). Knowledge about the major drivers of gene flow is important for the conservation and management of aquatic plants.

The rapid emergence and multidisciplinary nature of landscape genetics has introduced several hypotheses that assess the relationship between spatially explicit landscape variables and genetic measures (Manel and Holderegger, 2013; Hall and Beissinger, 2014). This is in an effort to understand whether dynamic landscape complexities limit gene flow thus resulting in discrete population structure (isolation by barrier, IBR), or clinal population structure is as a result of isolation by distance (IBD) and/or isolation by environment (IBE) (Meirmans, 2012; Landguth and Schwartz, 2014). Clustering of individuals in natural populations could be due to IBD, IBE, IBR or a combination of either (Cushman et al. 2006; Andrews, 2010; Wang and Bradburd, 2014; van Strien et al., 2015). IBD pattern results from geographical isolation, which limits gene flow and promotes genetic drift (Wright, 1943). This model assumes that genetic structure is driven by the linear relationship between geographic distance and genetic distance. It disregards the effects of nonlinear complex landscapes, environmental variables and historical processes that influence gene flow (Jenkins et al., 2010). IBR model considers landscape features and historical processes in explaining the impediment to gene flow (McRae, 2006). Both IBD and IBR are concerned with the limitations to the dispersal of gametes, but fails to accommodate the effects of environmental variables. Moreover, discrete population structure can be misinterpreted as an IBD or IBR pattern, and this may confound identification of environmental contributions resulting in overestimation of population clustering (Tucker et al., 2014). In contrast, IBE considers the contribution of environmental heterogeneity in shaping the distributions of spatial genetic variation independent of geographical isolation (Wang and Bradburd, 2014). Most empirical research on population genetics has focused on gene flow and genetic drift as drivers of genetic structure ignoring environmental-driven natural selection (Orsini et al., 2013). Understanding the environmental variables shaping genetic variation in natural populations is, therefore, important in evolutional studies and thus environmental gradients should be quantified and ecological approach included in population genetics studies (Lee and Mitchell-Olds, 2011). This information could be crucial in defining conservation and management units for the ecologically and economically important species (Segelbacher et al., 2010; Luque et al., 2012).

The wild rice *Zizania latifolia* is a perennial aquatic grass, and an important ecological and genetic resource in China. To date, the genetic differentiation of the wild rice across China has been attributed to IBD (Chen et al., 2017; Zhao et al., 2018; 2019). This is despite the fact that the distribution of wetlands in China, which represents the extant natural habitat for the species, has more unique features besides being expansive and patchy. The wetland ecosystem has a regional (Eastern) pattern with swamps distributed in the Northeast region and lakes in the middle-lower reaches of the Yangtze River and the Qinghai Tibet Plateau (Wang et al., 2012). Mangrove wetlands are distributed in the southeast coastal area. Topographically, the Eastern region, where most of the wetlands are found, is characterized by three plains; North, Northeast and Middle-Lower Yangtze plains amongst which foothills and hills are interspersed. This is the most economically important region characterized by a dense population, agriculture, and industries (An et al., 2007). China natural wetlands characteristics and distribution of *Z. latifolia*, therefore, presents an opportunity to study the landscape genetics of widespread species that have for long been neglected by conservation genetics community (Morente-López et al., 2018). This would aid in delineating the impact of island-like aquatic species distribution as well as human- and climate-change-induced habitat degradation (An et al., 2007) on the genetic structure of riparian plants at different spatial and environmental gradients. Further, the environmental gradient provides a perfect model for space-for-time substitution in assessing the long term aquatic ecosystem response to the changing environment (De Frenne et al., 2011).

In this study, we aimed to study the genetic diversity and population structure of 28 natural population of *Z. latifolia* based on 25 simple sequence repeat (SSR) markers and determine the influence of IBD, IBR and IBE in shaping the genetic structure. Based on the patchy nature of the wild rice habitat and the extensive latitudinal geographic range of the studied populations, it was predicted that *Z. latifolia* would show strong spatial genetic structure driven by diverse biotic and abiotic factors. It was expected that population decline and discontinuity, due to habitat degradation and fragmentation (An et al., 2007), could have considerable influence on wild rice reproduction system and gene flow, resulting in low genetic diversity. Further, we hypothesized that *Z. latifolia* genetic structure, in the expansive range and heterogeneous landscape, could be explained by combined effects of geographic isolation (IBD), gene-flow-limiting landscape variables (IBR) and differences in environmental variables between populations (IBE).

## 2 MATERIALS AND METHODS

### 2.1 Focal species, sampling and SSR data

The wild rice *Zizania latifolia*, commonly known as the Chinese wild rice, belongs to the family Poaceae, tribe Oryzeae. It shares the genus *Zizania* L. with other three species distributed in North America *(Z. aquatic Z. palustris* and *Z. texana*) (Xu et al., 2010). *Z. latifolia* is a perennial species distributed in Eastern Asia and well differentiated from the North American species (Brown, 1950; Dore, 1969; Chen et al., 1990; Kennard et al., 2000). It grows along the margins of lakes and rivers, ponds and marshes, and reproduces sexually by seeds or asexually by rhizomes (Xu and Zhou, 2017). In China, the wild rice is an important ecological species that is exploited for wastewater purification due to its high nutrient uptake capacity (Liu et al., 2007; Zhou et al., 2007; Peng et al., 2013) and high clonal reproduction (Chen et al., 2017). It also carries important genetic traits that include disease and pest resistance, elite grain, and tolerance to cold and flooding (Liu et al., 1999; Yu et al., 2006; Shen et al., 2011; Wang et al., 2013). Natural populations of *Z. latifolia* are distributed along the East of China along a wide stretch of latitudinal zones (21°N -50°N) (Wagutu et al., 2020). This region has five major eco-geographic regions that vary in biotic and abiotic factors that influence gene flow and local adaptation (Wu et al., 2003).

Twenty-eight populations of *Z. latifolia* were collected across China from Heilongjiang province to Guangdong province (Table 1) in a cluster sampling approach (Li et al., 2017). This was based on the size of the population, which should be large enough to allow a random selection of at least 20-23 samples at intervals of 10 meters. Samples were collected in the natural wetlands; rivers, lake shores and marshes during early autumn as the wild rice approaches maturity. GPS coordinates were recorded at the center of each collection transect and mapped using ArcGIS (Figure 1). The two uppermost leaves from each selected individuals were harvested, individually dried in silica gel and stored in the laboratory for DNA extraction. Total genomic DNA was extracted from 0.5g dried leaves using a modified CTAB protocol (Doyle and Doyle, 1987). Twenty five SSR markers were used in this study, including 20 markers developed for *Z. latifolia* (Quan et al., 2008; Wagutu et al., 2020), three SSR markers for *Oryza sativa* (Wang et al., 2015) and two markers developed for *Z. texana* (Richards et al., 2007) (Supplementary Table S1). PCR amplification was performed following the protocol by Quan et al. (2008) and PCR products separated on a 6% denaturing polyacrylamide gel. Fragments were visualized by silver staining and alleles scored in reference to a 25bp DNA ladder (Promega, Madison, WI, USA).

**Table 1.**
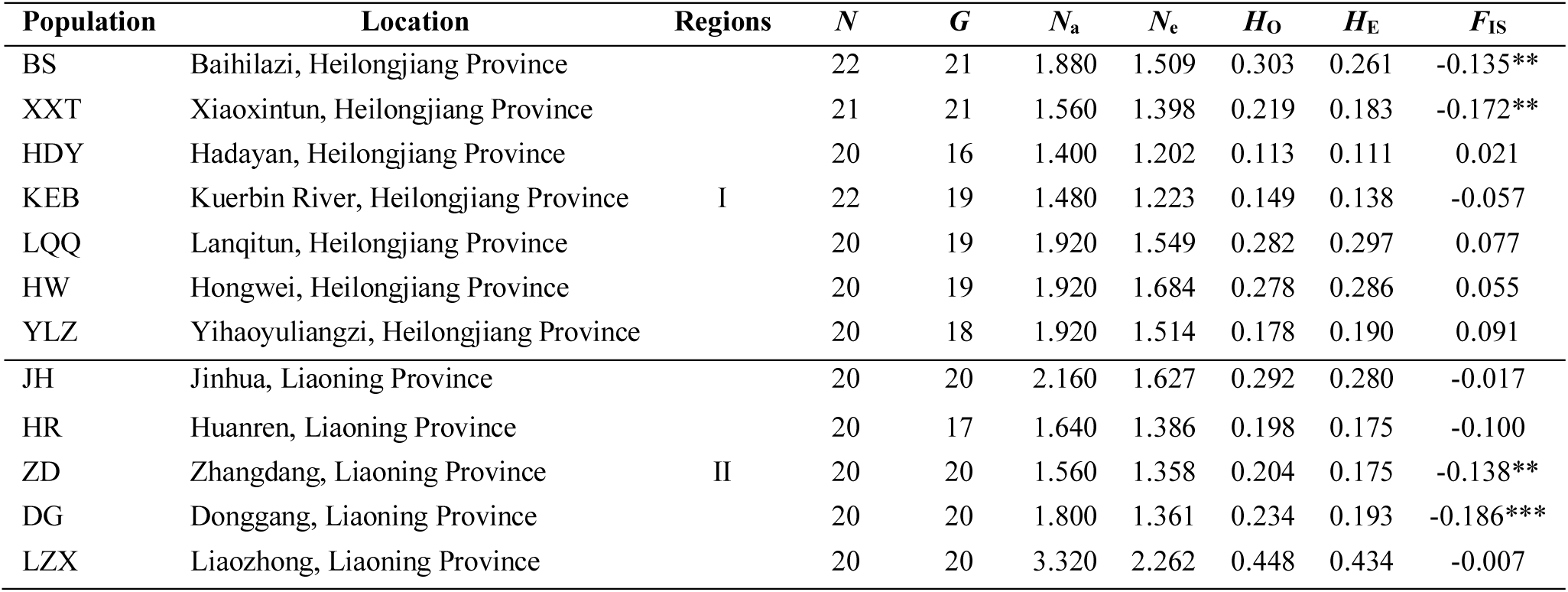
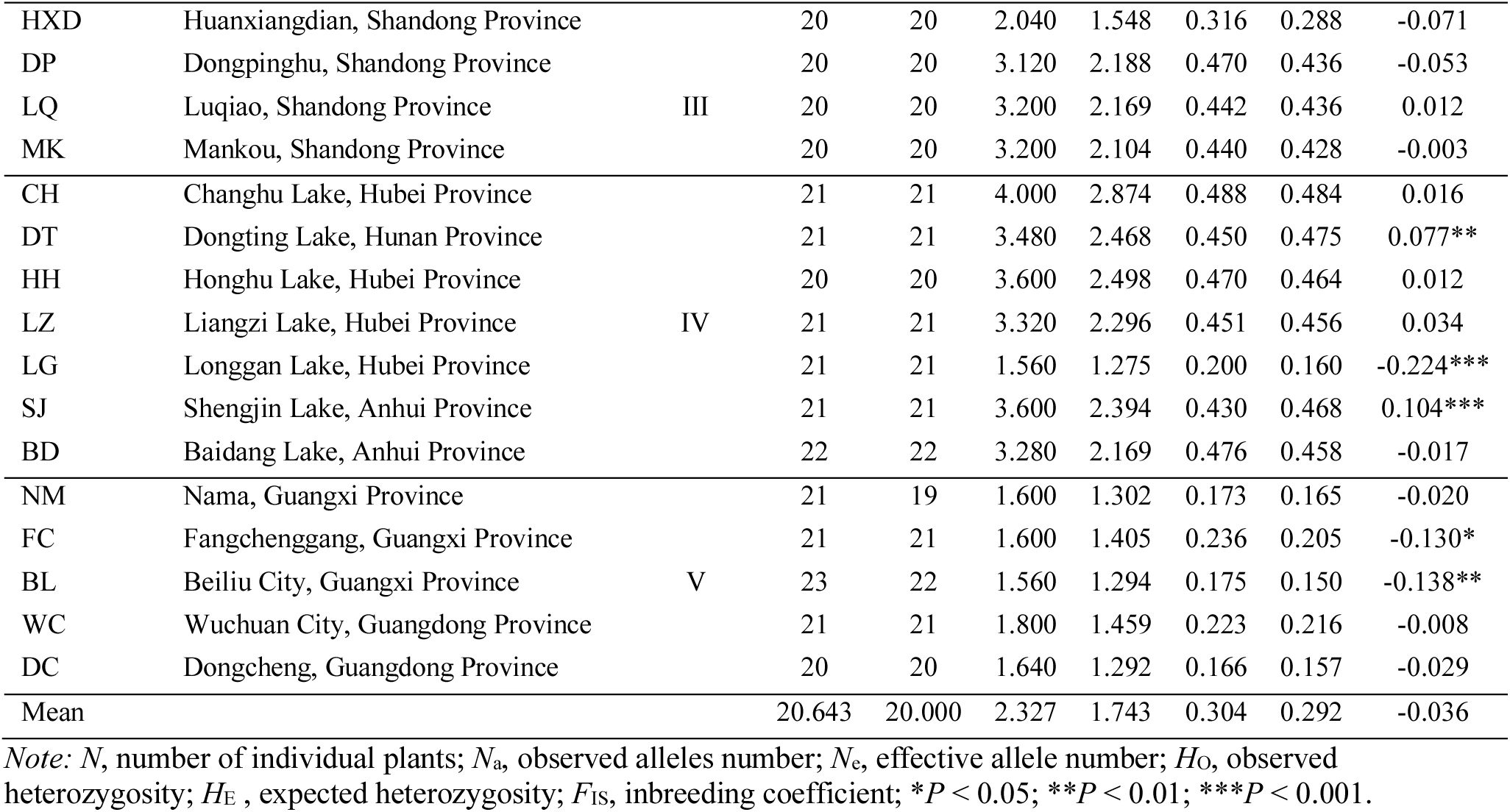
Geographical information and summary measures of clonal diversity and genetic variation for each of the 28 wild populations of *Zizania latifolia* along the five latitudinal regions starting from the north towards the south.

**Figure 1:**
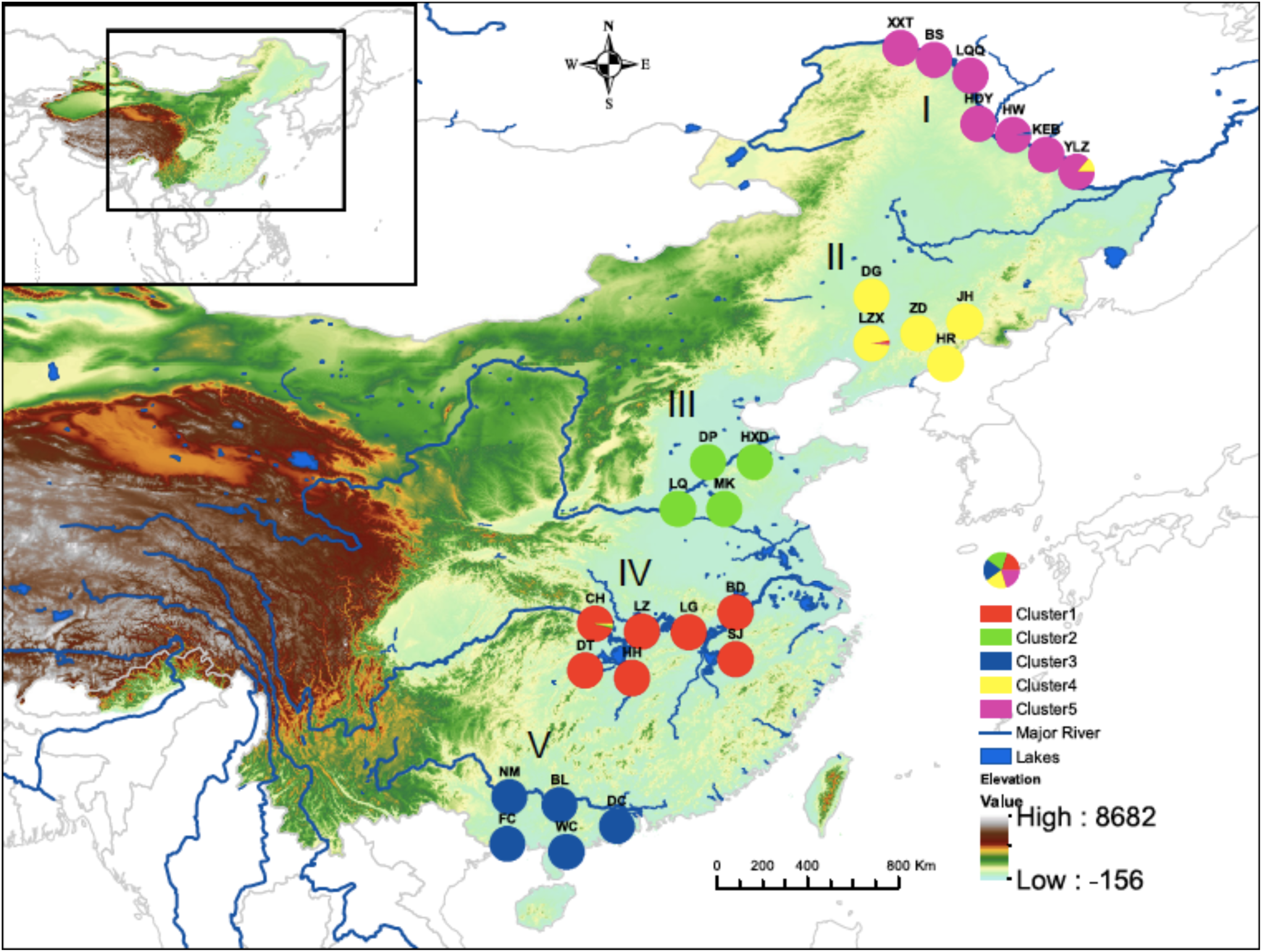
Geographical distribution of wild rice samples across the five (I to V) latitudinal regions in China and their respective genetic clusters each denoted by a different color.

GenoDive 2.0 (Meirmans et al., 2004) was used to identify clonal structure. At threshold zero, samples were assigned to their respective clones. The number of genotypes (*G*) was calculated and repeating genotypes excluded from further analysis. Micro-Checker 2.2.3 (Van Oosterhout et al., 2004) was used to test for null alleles and genotyping errors. Genetic diversity across all loci for every population and for each locus was estimated in terms of observed allele number (*N*_a_), the effective number of alleles (*N*_e_), and expected and observed heterozygosity (*H*_E_ and *H*_O_) using GenAlEx software (Peakall and Smouse, 2012). FSTAT (Goudet, 2001) was used to calculate the inbreeding coefficient (*F*_IS_), deviation from Hardy-Weinberg equilibrium and linkage disequilibrium.

### 2.2 Genetic structure among populations

FSTAT software was used to measure population genetic divergence by estimation of *F*_ST_ using 999 permutations for their significance. Based on *F*_ST_ values, an average level of gene flow (*Nm*) was estimated using the formula: [*Nm* = (1- *F*_ST_)/4*F*_ST_] (Slatkin and Barton, 1989). ARLEQUIN software (Schneider et al., 2000) was used to perform analysis of molecular variance (AMOVA) to determine genetic variation among and within populations. STRUCTURE program (Pritchard et al., 2000) was used to perform Bayesian clustering of the samples. Ten independent runs for each number of *K* clusters from one to ten was performed. A 20,000 iteration burn-in period followed by 100,000 Markov Chain Monte Carlo (MCMC) iterations were assumed for each run with correlated allele frequencies and admixture origin assumptions. To determine the value of *K*, the output was interpreted with Structure Harvester (Evanno et al., 2005; Earl and vonHoldt, 2012). However, Evanno’s *delta K* method has been reported to suffer philosophical and statistical errors (Verity and Nichols, 2016). Therefore, it was supplemented with Thermodynamic Integration (TI) method (Verity and Nichols, 2016). Here, *rmaverick* R package was used to estimate the true value of *K* by running 20 rungs for *K* = 1 to 10 with a burn-in period of 10,000 iterations followed by 50,000 MCMC iterations under the admixture model. The value of *K* was estimated as described by Verity and Nichols (2016). Since both software did not differ in the estimation of *K*, Structure Harvester output was visualized by DISTRUCT v. 1.1 (Rosenberg, 2004). Identified clusters were analyzed for molecular variations (AMOVA) using ARLEQUIN software. Geographical distribution of the different clusters identified was mapped using ArcGIS 10.0 (Esri, Redlands, CA, USA).

### 2.3 Geographic and environmental influence on the genetic structure

To determine the drivers of the observed genetic structure in *Z. latifolia*, we tested for isolation by distance (IBD), isolation by resistance (IBR) and isolation by environment (IBE) hypotheses. Four matrices were obtained in order to explore the three assumptions. Genetic distance was calculated as pairwise *F*_ST_ using FSTAT software, while the geographic distance matrix was based on Euclidean distance between populations calculated using GenAlEx. To linearize genetic and geographic distances relationship, both matrices were transformed: [*F*_ST_/(1- *F*_ST_)] and log(Euclidean distance), respectively (Rousset, 1997). Nineteen environmental variables were extracted for the studied sites from BioClim’s 30s resolution dataset (Busby, 1991) with GIS details using ArcGIS software (Supplementary Table S2). PAST ver. 3.24 (Hammer et al., 2001) was used to reduce climatic variables by principal component analysis (PCA) based on 20 variables (19 bioclimatic variables and elevation). The first principle components (PC) was used as it represented 99.32% of the variation in the data. Environmental distance matrix was calculated using the first principal component using *vegan* package in R (Oksanen et al., 2018). Lastly, resistance distance was calculated as the conductance matrix on a 30-arc seconds resolution digital elevation model (DEM) (Danielson and Gesch, 2011) of the study site using *gdistance* package implemented in R (van Etten, 2017). The resistance distance was obtained by calculating the effective resistance between populations when the DEM grid is conceived as an electrical circuit (McRae et al., 2008). Since wind dispersal is considered a major factor in *Zizania latifolia* gene flow, DEM was slope-modeled in ArcGIS 10.0. The grid cells with high values (high slope) contribute high resistance to gene flow and those with low value contribute the least resistance. Movement from any cell within the grid followed the standard eight directions (McRae et al., 2008).

To compete the three hypotheses, we used reciprocal causal modeling (RCM) (Cushman et al., 2006; 2013b) that eliminates the simple Mantel test’s (Mantel, 1967) limitations of spurious correlations and type I error (Cushman and Landguth, 2010). The RCM method directly compares the different hypotheses and identifies whether any of them is relatively supported independently of the other alternatives. The approach is based on a pair of partial Mantel tests derived from two models at a time. First, partial Mantel was performed between genetic distance (Gen) and IBD, partialling out IBR (Gen∼IBD|IBR). The second partial Mantel was performed between the genetic distance (Gen) and IBR, partialling out the IBD (Gen∼IBR|IBD). The difference between the two partial Mantel tests (IBD|IBR- IBR|IBD) was the relative support for IBD relative to IBR (Cushman et al., 2013a). In that case, the difference between the partial mantel should be positive if IBD is correct, and zero or negative if the IBR is correct. The full matrix of the partial Mantel tests differences between pairs of alternative hypotheses was computed. A hypotheses was regarded as fully supported, independent of all alternatives, if all the values in its column were positive and the values in the row were negative (Cushman et al., 2013a, b). Correlation values and significance values for the Mantel model combinations were calculated through 9999 corrected permutations using *vegan* R package.

We further evaluated the relationship between genetic diversity estimators and geographic and environmental variables through generalized linear models (GLMs) using PAST ver. 3.24 (Hammer et al., 2001) and spatially explicit mixed modeling (Morente-López et al., 2018) using *spaMM* R package (Rousset and Ferdy, 2014). Firstly, generalized linear models were used to explore the contribution of each environmental variable to genetic diversity estimators (*H*_O_, *H*_E_ and *N*_a_). Secondly, spatially explicit generalized linear mixed models (spatial GLMMs) were developed using genetic, geographic (coordinates and elevation) and environmental data. Here, we used genetic diversity estimators (*H*_O_, *H*_E_ and *N*_a_) as response variables, each of the 19 environmental variables as fixed effects and geographical coordinates and elevation as random effects. We transformed environmental variables as required, including using their squared values to account for non-linearity. Full models (e.g. [*F*_IS_ ∼ bio_1 + (1|lat+long/Elevation)]) and null models (e.g. [*F*_IS_ ∼ 1 + (1|lat+long/Elevation)]) were tested for associated likelihood ratio to obtain the *P*-value.

## 3 RESULTS

From 578 individuals, 570 multilocus genotypes were identified, and no genotype was shared among populations. In 19 populations, every individual was of unique genotype (Table 1). We did not find evidence of null alleles or genotyping error using Micro-Checker. At the population level, the range of observed heterozygosity (*H*_O_) and expected heterozygosity (*H*_E_) were 0.113 to 0.488 and 0.111 to 0.484, respectively (Table 1). Fixation index ranged from *F*_IS_ = -0.334 to *F*_IS_ = 0.104 with a mean of -0.036. Deviation from HWE was detected in 257 out of the 700 locus-population combinations (36.7%) at *P* = 0.05, which is expected for natural population at a broad geographical scale (Garnier-Géré and Chikhi, 2013). Significant genotypic disequilibrium was detected in 86 out of 300 locus pairs (*P* = 0.001), but after Bonferroni correction, none of the locus pairs was in significant genotypic disequilibrium. This indicated that loci were unlinked and statistically independent of each other.

### 3.1 Genetic structure and differentiation among populations

The genetic differentiation across all populations based on *F*_ST_ was 0.579 (Table 2). Pairwise comparisons of *F*_ST_ were significant for genetic differentiation between populations (*P* = 0.01). Moreover, the levels of differentiation were high; *F*_ST_ = 0.103 to 0.823 (Supplementary Table S3). AMOVA showed high population differentiation at Ø_pt_ = 0.578 (*P*=0.000). Variation among populations was 58%, while 42% of the variation was within population, both statistically significant (*P* = 0.001) (Table 3). Bayesian clustering using structure identified optimum *K*=5 using the Evanno’s delta *K* as well as TI method (Pritchard et al., 2000; Verity and Nichols, 2016). Populations from the five regions clustered according to their respective collection sites (Figure 2). Membership proportion of predefined populations in each of the 5 clusters ranged from 0.857 to 0.997. AMOVA showed that 36% of variation was among clusters (Table 3). The five identified clusters were mapped using ArcGIS (Figure 1).

**Table 2.** Genetic diversity found at 25 microsatellite loci in *Zizania latifolia*

| Locus | $A$ | $H_o$ | $H_E$ | $F_{ST}$ | $F_{IS}$ | $Nm$ |
| --- | --- | --- | --- | --- | --- | --- |
| ZM4 | 7 | 0.278 | 0.304 | 0.602 | 0.111 | 0.060 |
| ZM25 | 6 | 0.192 | 0.183 | 0.762 | -0.022 | 0.045 |
| ZM26 | 9 | 0.281 | 0.278 | 0.641 | 0.016 | 0.058 |
| ZM28 | 4 | 0.114 | 0.114 | 0.818 | 0.026 | 0.037 |
| ZM35 | 12 | 0.459 | 0.463 | 0.451 | 0.034 | 0.062 |
| ZM44 | 11 | 0.495 | 0.510 | 0.386 | 0.050 | 0.059 |
| RM14233 | 12 | 0.379 | 0.339 | 0.599 | -0.090 | 0.060 |
| RM20118 | 6 | 0.410 | 0.342 | 0.556 | -0.170 | 0.062 |
| RM28090 | 7 | 0.529 | 0.435 | 0.431 | -0.197 | 0.061 |
| Zt1 | 7 | 0.290 | 0.250 | 0.692 | -0.130 | 0.053 |
| Zt23 | 6 | 0.187 | 0.187 | 0.718 | 0.027 | 0.051 |
| ZL1 | 30 | 0.706 | 0.607 | 0.292 | -0.136 | 0.052 |
| ZL3 | 29 | 0.413 | 0.480 | 0.476 | 0.167 | 0.062 |
| ZL4 | 28 | 0.396 | 0.444 | 0.512 | 0.131 | 0.062 |
| ZL5 | 28 | 0.416 | 0.454 | 0.511 | 0.108 | 0.062 |
| ZL9 | 5 | 0.211 | 0.212 | 0.560 | 0.095 | 0.062 |
| ZL10 | 7 | 0.276 | 0.277 | 0.510 | 0.028 | 0.062 |
| ZL31 | 7 | 0.346 | 0.305 | 0.427 | -0.109 | 0.061 |
| ZL32 | 8 | 0.315 | 0.291 | 0.482 | -0.056 | 0.062 |
| ZL36 | 5 | 0.213 | 0.193 | 0.739 | -0.085 | 0.048 |
| ZL42 | 5 | 0.105 | 0.110 | 0.445 | 0.079 | 0.062 |
| ZL43 | 4 | 0.117 | 0.115 | 0.432 | 0.014 | 0.061 |
| ZL55 | 4 | 0.072 | 0.072 | 0.875 | 0.016 | 0.027 |
| ZL56 | 5 | 0.344 | 0.273 | 0.649 | -0.236 | 0.057 |
| ZL57 | 4 | 0.067 | 0.059 | 0.908 | -0.125 | 0.021 |
| Mean | 10.240 | 0.304 | 0.292 | 0.579 | -0.016 | 0.061 |

Note: $A$ , average number of alleles per locus; $H_E$ , expected heterozygosity; $H_o$ , observed heterozygosity; $F_{ST}$ , coefficient of genetic differentiation; $Nm$ , gene flow

**Table 3.**
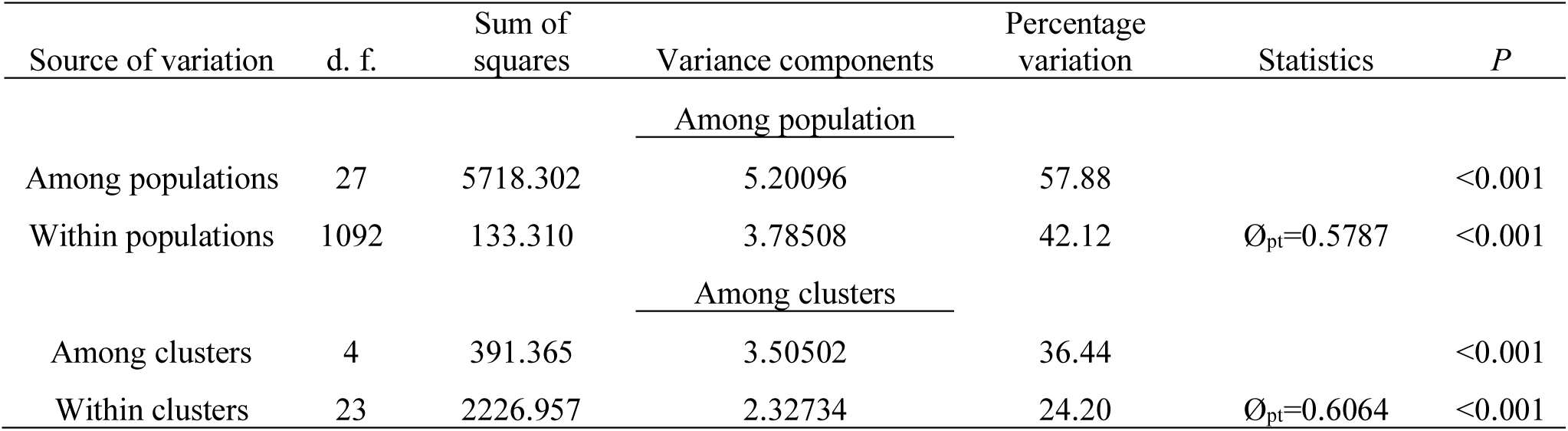
AMOVA design and results (average over 25 loci)

| Source of variation | d. f. | Sum of squares | Variance components | Percentage variation | Statistics | *P* |
| --- | --- | --- | --- | --- | --- | --- |
| <u>Among population</u> |  |  |  |  |  |  |
| Among populations | 27 | 5718.302 | 5.20096 | 57.88 |  | <0.001 |
| Within populations | 1092 | 133.310 | 3.78508 | 42.12 | $\phi_{pt}=0.5787$ | <0.001 |
| <u>Among clusters</u> |  |  |  |  |  |  |
| Among clusters | 4 | 391.365 | 3.50502 | 36.44 |  | <0.001 |
| Within clusters | 23 | 2226.957 | 2.32734 | 24.20 | $\phi_{pt}=0.6064$ | <0.001 |

**Figure 2:**
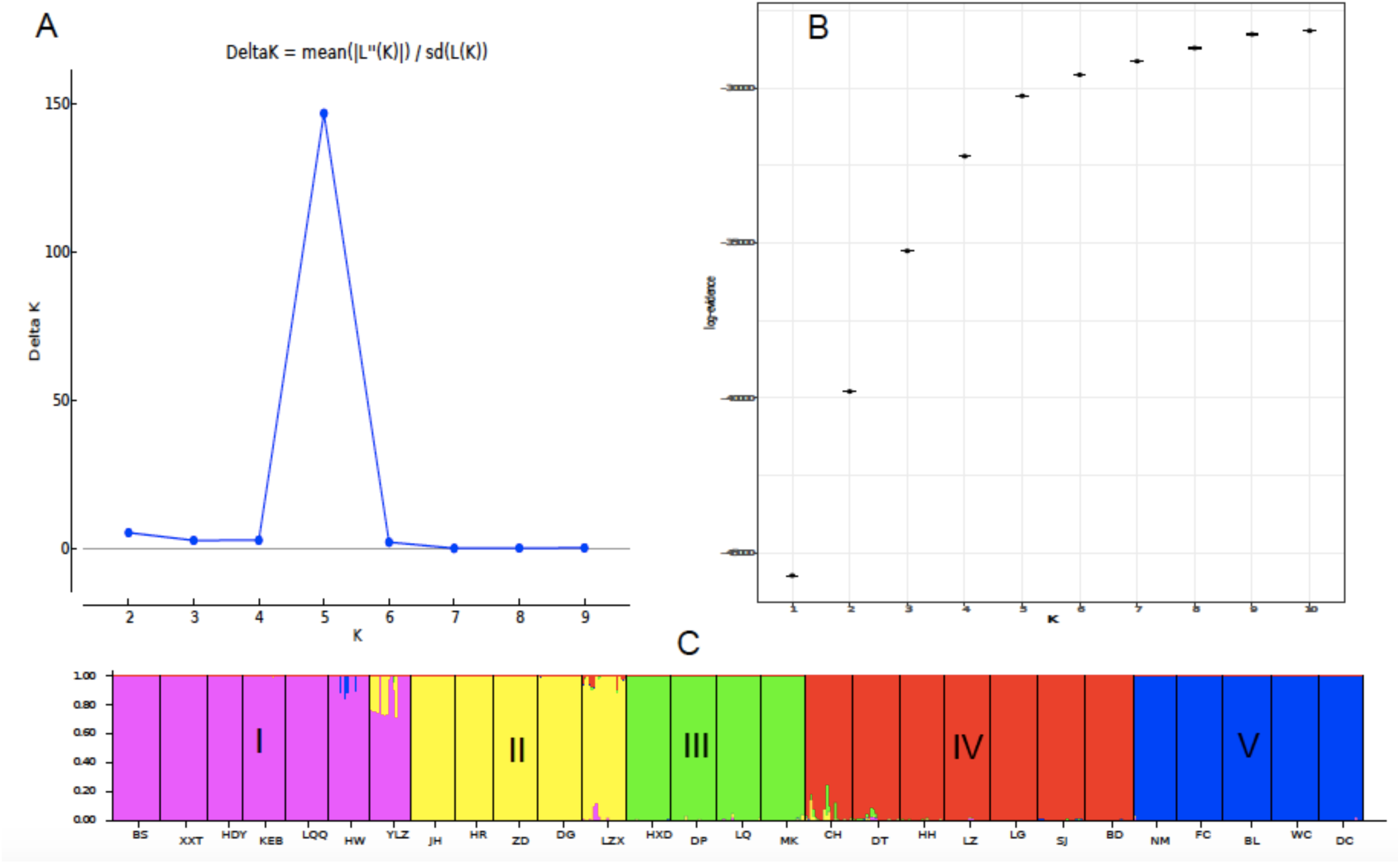
Bayesian analysis of population structure (STRUCTURE) for *K* = 5 inferred by DISTRUCT as calculated by: **(A)**, Evanno’s delta *K* and; **(B)**, log-evidence in thermodynamic integration. **(C)** STRUCTURE composition of individuals, with each of the clusters corresponding to the five latitudinal regions and color-coded similar to Figure 1.

**Figure 3:**
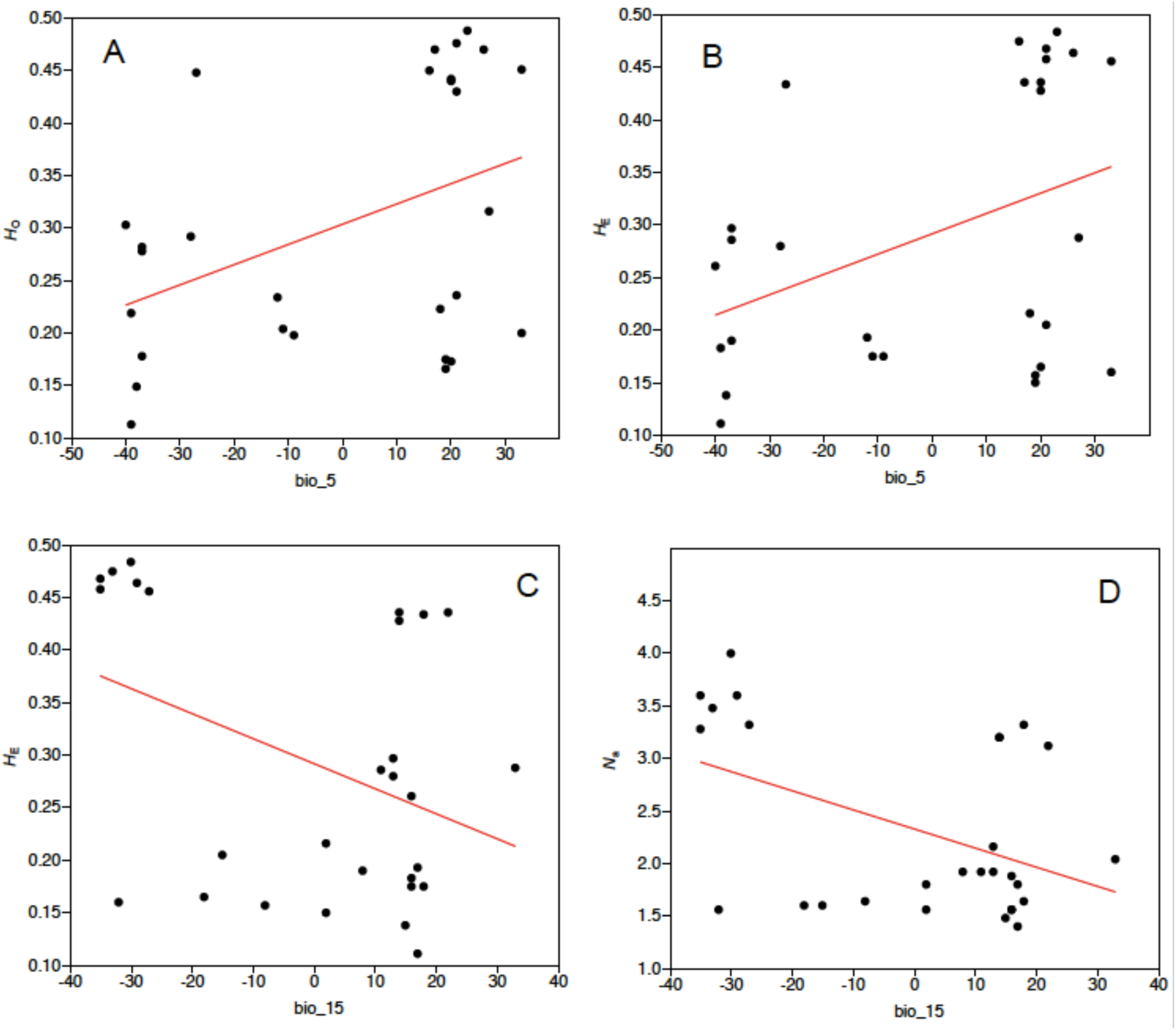
Significant relationships between genetic diversity estimators and the two most important environmental variables, based on variance inflation factor analysis. **(A)** *H*_O_, observed heterozygosity; **(B and C)** *H*_E_, expected heterozygosity; **(D)** *N*_a_, number of alleles per population. bio_5; Maximum temperature of warmest month, bio_15; precipitation seasonality.

### 3.2 Geographic and environmental influence on the genetic structure

IBE model was fully supported based on the relative support values of the reciprocal causal model. The column had positive values and the row had negative values (Table 4) indicating that it explained the genetic structure independent of the alternative hypotheses. Moreover, the partial Mantel tests between genetic distance and environmental distance controlling for both resistance and geopgraphy showed significant positive correlations (Table 4). IBR model had one positive value in the column when competed with IBD and partial Mantel tests were all significant. IBD model had all negative value in the column and thus had the least relative support. However, partial Mantel tests when controlling for IBR showed a significant positive correlation. In simple Mantel, environmental distance had the highest significant positive correlation with genetic distance followed by resistance distance and then geographical distance (Table 4).

**Table 4.** Reciprocal causal modeling, partial and simple Mantel results for IBD, IBR and IBE

|  |  | IBD | IBR | IBE |
| --- | --- | --- | --- | --- |
|  |  | Geo | Res | Env |
| I: Reciprocal causal modeling matrix |  |  |  |  |
| IBD | Geo | 0 | 0.0851 | **0.4961** |
| IBR | Res | -0.0851 | 0 | **0.0392** |
| IBE | Env | **-0.4961** | **-0.0392** | **0** |
| II: Simple and Partial Mantel correlation matrix |  |  |  |  |
| IBD | Geo | 0.5830*** | 0.2684*** | 0.3063*** |
| IBR | Res | 0.1833** | 0.6051*** | 0.2724*** |
| IBE | Env | -0.1898 | 0.2332*** | 0.6160*** |

Note: IBD, isolation by distance; IBR, isolation by resistance; IBE, isolation by environment; Geo, geographic distance; Res, resistance distance; Env, environmental distance.
(I) Reciprocal causal modeling matrix; columns indicate test model and rows indicate alternative models. Each value represents the relative support of the test model
(II) Simple and Partial Mantel correlation matrix. Columns indicates test model and rows indicate partialled-out models. Values are r values for correlations, diagonal values are the simple Mantel test r of a variable. \* $P < 0.05$ ; \*\* $P < 0.01$ ; \*\*\* $P < 0.001$ .

Generalized linear models showed that seven of the 19 bioclimatic variables had significant contribution to genetic diversity of *Z. latifolia* (Table 5). However, spatially explicit mixed models with coordinates as random effect showed no significant influence of environmental variables to genetic diversity. When elevation was used as the random effect in the mixed models, bio_15 showed significant influence (*P* = 0.038) on number of alleles (*N*_a_). We tested the seven environmental variables showing significant contribution to genetic diversity in GLM for collinearity using variance inflation factor (VIF) analysis (Helsen et al., 2017). Two variables; bio_5 and bio_15 had VIF value below 2 and were therefore identified as the best environmental variables responsible for the genetic diversity of *Z. latifolia*.

**Table 5.** Generalized linear model for the influence of environmental variables on genetic diversity measures. Values are significant at *P*-values ≤ 0.05

| Independent variables | Dependent variables |  |  |
| --- | --- | --- | --- |
| | $H_E$ | $H_O$ | $N_a$ |
| bio_4 | – | – | 0.012 |
| **bio_5** | **0.024** | **0.017** | – |
| bio_10 | – | 0.046 | 0.030 |
| bio_14 | – | – | 0.032 |
| **bio_15** | **0.032** | – | **0.0090** |
| bio_17 | – | – | 0.020 |
| bio_19 | – | – | 0.043 |

## 4 DISCUSSION

In this study, *Zizania latifolia* showed relatively low genetic diversity (*H*_E_ = 0.292). A similar level of genetic diversity was reported in its natural populations from the Northeast wetland (*H*_E_ = 0.328; Chen et al., 2017) and the lower-middle Yangtze wetland (*H*_E_ = 0.271; Wang et al., 2015) using SSR markers. Our results are consistent with the expected average genetic variability for the family Poaceae (*H*_E_ = 0.201), which is estimated based on characteristics that influence gene flow (Hamrick and Godt, 1996). These results also correspond to Northern American wild rice *Z. palustris* genetic diversity (*H*_E_ = 0.236; Lu et al., 2005). Further, *Z. latifolia* shows lower genetic diversity compared to *Z. texana* (*H*_E_ = 0.662; Richards et al., 2007). However, the two studies employed different sampling approaches, with Richards et al. (2007) considering a single hydrologically connected river system, while we sampled different populations across large geographical and environmental gradient (Figure 1). A higher than average genetic diversity in populations in region III, along the Yangtze River (*H*_E_ = 0.424) and region IV, along the Yellow River (*H*_E_ = 0.426) was observed. Similar results were reported by Chen et al. (2012) and Zhao et al. (2018) who showed that *Z. latifolia* wild populations in the middle and lower reaches of the Yangtze basin exhibit high genetic diversity. This could be explained that, the central and southeast China wetlands are characterized by episodic floods that create temporary hydrological connectivity (Wang et al., 2008) resulting in gamete exchange between population. Genetic diversity is then maintained by the high clonal capacity of the species.

### 4.1 Genetic structure and differentiation among populations

High level of genetic differentiation among populations was detected (*F*_ST_ = 0.579 *D*_est_ = 0.598). Our results are comparable to *Z. latifolia* genetic differentiation reported recently (*F*_ST_ = 0.481, Xu et al., 2008; *F*_ST_ = 0.405, Chen et al., 2017). AMOVA analysis also showed high population differentiation at Ø_pt_ = 0.578 (*P*=0.001), with 58% variation attributed to between populations and 42% to within population (*P* = 0.001) variation. This was also comparable to *Z. latifolia* population differentiation (Ø_pt_ = 0.598) reported by Xu et al. (2015). Moreover, high population differentiation was also reported in the North American relatives of *Z. latifolia* (*Z. aquatica*: *F*_ST_ = 0.607; *Z. palustris*: *F*_ST_ = 0.468; Xu et al., 2015). The observed high genetic divergence is attributable to decreased gene flow between populations (*Nm* = 0.061). According to Wright (1969), when *Nm* is perceived as the exchange of migrants between demes per generation, a value > 1 result in little divergence, while value < 1 results in increased divergence. In our case, conventional fragmentation of wetlands into islands within the expansive terrestrial habitat and induced fragmentation could explain the negligible gene flow and the observed high genetic differentiation (Wagutu et al., 2020). Moreover, inbreeding system detected in some populations could have contributed to the high genetic differentiation. Bayesian clustering using STRUCTURE software and *rmaverick* (R package) identified five clusters (Figure 2). All the five clusters corresponded to the wetland from which the samples were collected. The clustering pattern can be attributed to the *Z. latifolia* habitat, which is discrete and patchy within an expansive geographical range (Zhao et al., 2018). Further, dispersal of its gametes and somatic propagules depend on hydrological connectivity and landscape features. In such spatially isolated populations under a heterogenous landscape, wind and water dispersal is most effective locally creating locally identical genotypes. Environmental difference between the geographically isolated populations promote phenotypic plasticity and genetic changes.

### 4.2 Environmental, landscape and geographical influence on genetic structure and genetic diversity

For the first time, this study tested for IBE and IBR patterns, besides the commonly tested IBD pattern in the natural populations of *Z. latifolia*. Interestingly, we found exclusive relative support by RCM for IBE pattern, partial support for IBR, but not IBD, in shaping genetic structuring of the species. Similarly, Šurinová et al. (2019) reported that environmental variables, but not geographical distance, explained genetic relatedness among a perennial grass species (*Festuca rubra*) populations. Wu et al. (2019), when comparing IBD and IBE in aquatic species *Ranunculus subrigidus* on the Qinghai-Tibetan plateau, also found exclusive support for environmental isolation. Further, IBE has been reported as a more common phenomenon compared to IBD in a wide range of taxa (Shafer et al., 2013; Sexton et al., 2014), although it has for long been neglected in population genetic studies. IBE can be generated by various ecological processes that include; natural and sexual selection against maladapted immigrants, poor hybrid fitness and biased dispersal (Wang and Bradburd, 2014). In natural selection, populations evolve traits suitable for the environment and thus acquire higher fitness compared to immigrants from different environments. Sexual selection and dispersal bias may occur as a result of phenotypic plasticity. Here, environmental conditions influence reproductive physiology affecting synchronization of mating and dispersal processes such flowering and gamete release. Moreover, if immigrates survive long enough and are able to mate locally, hybrids would undergo selection against immigrant alleles (Sexton et al., 2014). The sexual reproduction of *Z. latifolia* could be one characteristic that makes natural and sexual selection as well as dispersal bias drive local adaptation and thus the observed pattern of IBE. However, the species also reproduces asexually through rhizomes and thus hybrids survival could be a tradeoff. The net effect of ecological processes targeting sexual reproduction and asexual reproduction, however, seems to favor local adaptation and hence the exclusive support of IBE by RCM.

In this study, IBE was further supported by the significant relationship between genetic diversity estimators and a number of environmental variables in the GLMs, although limited relationship in the spatial GLMMs (Table 5). Maximum temperature of the warmest month (bio_5) and precipitation seasonality (bio_15) were identified as the best environmental variables responsible for observed genetic differentiation based on VIF analysis. Temperature and precipitation are major drivers of plant adaptability through their influence on physiology and diversity (Hoffmann and Sgrò, 2011; Manel et al., 2012), forcing genetic change events and subsequent selection. Besides our study, influence of temperature and precipitation on genetic differentiation of both aquatic and terrestrial plants have been reported (Wang et al., 2016; Münzbergová et al., 2017). A significant relationship between precipitation seasonality (bio_15) and *N*_a_ was detected in our study. This indicates that the local adaptation, driven by interaction between precipitation and temperature, is responsible for *Z. latifolia* differentiation along the eastern China aquatic ecosystem. This could be due to the fact that the species is tolerant to a wide range of temperatures (Guo et al., 2007), but since it’s an aquatic species, the differences in precipitation along the sampling sites and between seasons has profound influence. As a perennial and aquatic species, *Z. latifolia* reproduction and growth will be affected by temperature and precipitation fluctuations. Flowering, vegetative growth and reproductive success of plants have been reported to respond to temperature changes (De FRENNE et al., 2011). Similarly, asynchronous flowering due to heterogenous rainfall patterns have been associated with IBE pattern (Garot et al., 2019).

IBR was the second best model explaining *Z. latifolia* genetic structuring. In partial Mantel, IBR had a positive correlation with genetic diversity when controlling for both geographical distance and environment. The model was partially supported by RCM (Table 4). Similarly, significant effect of cost distance on genetic differentiation has been reported in alpine ecosystem (Morente-López et al., 2018). Wetlands along eastern China are islands within terrestrial ecosystem interspersed with foothills and hills (An et al., 2007; Santamaria, 2002). Such natural barriers are important factors in the gene flow of flowering plants. Similar, albeit artificial barriers, have been reported to drive genetic differentiation in species that depend on wind, insects and birds for pollination and seed dispersal (Su et al., 2003). In the case of *Z. latifolia*, wind pollination seems to be inhibited by the complex landscape along the eastern China, and seed dispersal by lack of hydrological connectivity, which leads to genetic drift and thus population differentiation.

The exclusive IBE pattern detected contradicts Zhao et al. (2018; 2019) who reported a pattern of IBD along the eastern China populations of *Z. latifolia*. However, in their study, only IBD was tested using simple Mantel, for which we also found a strong positive correlation (*r*=0.5830; *P*=0.001). The observed direct correlation of genetic differentiation with geographical distance between populations could be due to the discrete pattern of *Z. latifolia* populations along the expansive geographical area. Moreover, rapid fragmentation and reclamation of the wetlands (An et al., 2007; Zhao et al., 2018) has increased population isolation, which inhibits wind- and water-mediated gamete dispersal. In our study, IBD was the least supported hypothesis in partial Mantel. When controlling for IBR, significant correlation between IBD and genetic differentiation was detected (Table 4). This could be because calculation of cost distance on DEM has an aspect of distance (Morente-López et al., 2018).

### 4.3 Implications for conservation and management

The genetic variation in the extant natural populations of *Z. latifolia* is relatively low and the high genetic divergence is shaped by environmental variation between groups of populations. The heterogenous landscape, discrete distribution of the natural populations coupled with wetland fragmentation and degradation has increased the isolation resulting in genetic isolation. Genetic diversity measures were sensitive to temperature and precipitation. Between 1956 and 2005, earth surface temperature has increased at an average of 0.65°C and might increase with further 1.8°C to 4°C by 2099 (IPCC, 2007). This is expected to affect the precipitation and together impact on species diversity and survival. Therefore, further extensive genetic and morphological studies need to be carried out to identify the role of the environment in shaping local adaptation of the *Z. latifolia*, besides the aquatic ecosystem response assessment. The use of a common garden approach in combination with genome-wide association studies (GWAS) could help elucidate patterns of local adaptation (de Villemereuil et al., 2016).

Our results show that gene flow is critically inhibited among the populations. Therefore, management and conservation approaches should include both *in-situ* and *ex-situ* considerations. Populations in the central and southeast China wetland (e.g. CH, LZX, and DT) that showed the highest genetic variability should be prioritized for germplasm collection and conservation. The genetic profiles of the sampled populations within each of the five clusters could constitute valuable traits that require conservation as well. Geographic barriers showed positive correlation to genetic structuring of *Z. latifolia*. This is related to topographical complexity, which impedes pollen dispersal. While nothing can be done about the topography, seed and somatic propagule dispersal through water can be enhanced. Where landscapes are disconnected, revegetation with broad genotypes can be used to promote gene flow, which may also include deliberate movement of propagules across fragmented landscapes (Reviewed by Sexton et al., 2014). Chen et al. (2017) proposed dredging the watercourses to achieve hydrological connectivity within each wetland. Populations along the Yangtze and Yellow river exchange gametes through periodical floods, which shows that this approach is feasible in other wetlands.

## 5 Data accessibility

All datasets for this study are included in the article’s Supplementary Material

## 6 Authors’ contribution

YC conceived the idea and designed the research project. WL gave suggestions to the design of the study. YC, XF and WF collected the samples and assembled experiment materials. GKW, XF performed the experiment. GKW analyzed the data and wrote the manuscript. All authors contributed to the revision and final editing of the manuscript prior to submission.

## 7 Funding

This work was supported by Chinese Academy of Sciences (Y855291B01) and the CAS-TWAS President’s Fellowship for International Doctoral Students.

## 8 Supplementary material

**Supplementary Table S1**. Details of SSR primer sequences used in this study

**Supplementary Table S2**. Genetic diversity measures, geographical variables and bioclimatic variables used in in this study

**Supplementary Table S3**. *F*_ST_ pairwise matrix, all significant at *P* = 0.05

## Notes

### Competing Interest Statement

The authors have declared no competing interest.

## References

An, S., Li, H., Guan, B., Zhou, C., Wang, Z., Deng, Z., et al. (2007). China’s natural wetlands: Past problems, current status, and future challenges. AMBIO: A Journal of the Human Environment 36(4), 335–342. https://doi.org/10.1579/0044-7447(2007)36[335:CNWPPC]2.0.CO;2

Andrews, C. A. (2010). Natural selection, genetic drift, and gene flow do not act in isolation in natural populations. Nature Education Knowledge 3(10), 5.

Barrett, S. C. H., Eckert, C. G., and Husband, B. C. (1993). Evolutionary processes in aquatic plant populations. Aquatic Botany 44(2–3), 105–145. https://doi.org/10.1016/0304-3770(93)90068-8

Bell, G., and Collins, S. (2008). Adaptation, extinction and global change. Evolutionary applications 1(1), 3–16. https://doi.org/10.1111/j.1752-4571.2007.00011.x

Brown, W. V. (1950). A Cytological Study of Some Texas Gramineae. Bulletin of the Torrey Botanical Club 77(2), 63. https://doi.org/10.2307/2482267

Busby, J. (1991). BIOCLIM – A bioclimate analysis and prediction system. Plant Protection Quarterly 6, 8–9.

Catullo, R. A., Llewelyn, J., Phillips, B. L., and Moritz, C. C. (2019). The Potential for Rapid Evolution under Anthropogenic Climate Change. Current Biology 29(19), R996–R1007. https://doi.org/10.1016/j.cub.2019.08.028

Cao, Y., Neif, É., Li, W., Coppens, J., Filiz, N., Lauridsen, T., et al. (2015). Heat wave effects on biomass and vegetative growth of macrophytes after long-term adaptation to different temperatures: A mesocosm study. Climate Research 66(3), 265–274. https://doi.org/10.3354/cr01352

Chen, S. L., Qin, H. Z., Jin, Y. X., and Shu, P. (1990). Preliminary studies on systematic position and evolution of Zizania L. (Gramineae). In: Proc. Intern. Symp. Bot. Gard. Nanjing pp. 593–605.

Chen, Y.-Y., Chu, H.-J., Liu, H., and Liu, Y.-L. (2012). Abundant genetic diversity of the wild rice *Zizania latifolia* in central China revealed by microsatellites. Annals of Applied Biology 161(2), 192–201. https://doi.org/10.1111/j.1744-7348.2012.00564.x

Chen, Y., Liu, Y., Fan, X., Li, W., and Liu, Y. (2017). Landscape-Scale Genetic Structure of Wild Rice *Zizania latifolia*: The Roles of Rivers, Mountains and Fragmentation. Frontiers in Ecology and Evolution 5. https://doi.org/10.3389/fevo.2017.00017

Cushman, S. A., and Landguth, E. L. (2010). Spurious correlations and inference in landscape genetics. Molecular Ecology 19(17), 3592–3602. https://doi.org/10.1111/j.1365-294X.2010.04656.x

Cushman, S. A., McKelvey, K. S., Hayden, J., and Schwartz, M. K. (2006). Gene flow in complex landscapes: Testing multiple hypotheses with causal modeling. The American Naturalist 168 (4), 486–499. https://doi.org/10.1086/506976

Cushman, S. A., Shirk, A. J., and Landguth, E. L. (2013a). Landscape genetics and limiting factors. Conservation Genetics 14(2), 263–274. https://doi.org/10.1007/s10592-012-0396-0

Cushman, S., Wasserman, T., Landguth, E., and Shirk, A. (2013b). Re-Evaluating Causal Modeling with Mantel Tests in Landscape Genetics. Diversity 5(1), 51–72. https://doi.org/10.3390/d5010051

Danielson, J. J. and Gesch, D. B. (2011). Global multi-resolution terrain elevation data 2010 (GMTED2010): U.S. Geological Survey Open-File Report 26, 2011–1073.

Davidson, N. C. (2014). How much wetland has the world lost? Long-term and recent trends in global wetland area. Marine and Freshwater Research 65(10), 934–941. https://doi.org/10.1071/MF14173

De Frenne, P., Brunet, J., Shevtsova, A., Kolb, A., Graae, B. J., Chabrerie, O., et al. (2011). Temperature effects on forest herbs assessed by warming and transplant experiments along a latitudinal gradient. Global Change Biology 17(10), 3240–3253. https://doi.org/10.1111/j.1365-2486.2011.02449.x

de Villemereuil, P., Gaggiotti, O. E., Mouterde, M., and Till-Bottraud, I. (2016). Common garden experiments in the genomic era: New perspectives and opportunities. Heredity 116(3), 249–254. https://doi.org/10.1038/hdy.2015.93

Dore, W.G. (1969). Wild rice. Can. Dept. Agric. Publ. no. 1393, Ottawa, Ontario, Canada.

Doyle, J. J., and Doyle, J. L. (1987). A rapid DNA isolation procedure for small quantities of fresh leaf tissue. Phytochem. Bull. 19, 11–15.

Earl, D. A., and vonHoldt, B. M. (2012). STRUCTURE HARVESTER: A website and program for visualizing STRUCTURE output and implementing the Evanno method. Conservation Genetics Resources 4(2), 359–361. https://doi.org/10.1007/s12686-011-9548-7

Evanno, G., Regnaut, S., and Goudet, J. (2005). Detecting the number of clusters of individuals using the software structure: A simulation study. Molecular Ecology 14(8), 2611–2620. https://doi.org/10.1111/j.1365-294X.2005.02553.x

Garnier-Géré, P., and Chikhi, L. (2013). Population Subdivision, Hardy-Weinberg Equilibrium and the Wahlund Effect. In John Wiley & Sons Ltd (Ed.), ELS (p. a0005446.pub3). John Wiley & Sons, Ltd. https://doi.org/10.1002/9780470015902.a0005446.pub3

Garot, E., Joët, T., Combes, M.-C., and Lashermes, P. (2019). Genetic diversity and population divergences of an indigenous tree (*Coffea mauritiana*) in Reunion Island: Role of climatic and geographical factors. Heredity 122(6), 833–847. https://doi.org/10.1038/s41437-018-0168-9

Goudet, J. (2001). FSTAT ver 2.9.3, A program to Estimate and Test Gene Diversities and Fixation Indices. Available online at: https://www2.unil.ch/popgen/softwares/fstat.htm

Grant, P. R., Grant, B. R., Huey, R. B., Johnson, M., Knoll, A. H., and Schmitt, J. (2017). Evolution caused by extreme events. Philosophical transactions of the Royal Society of London. Series B, Biological sciences 372(1723), 20160146. https://doi.org/10.1098/rstb.2016.0146

Guo, H. B., Li, S. M., Peng, J., and Ke, W. D. (2007). *Zizania latifolia* Turcz. Cultivated in China. Genetic Resources and Crop Evolution 54(6), 1211–1217. https://doi.org/10.1007/s10722-006-9102-8

Hall, L. A., and Beissinger, S. R. (2014). A practical toolbox for design and analysis of landscape genetics studies. Landscape Ecology 29(9), 1487–1504. https://doi.org/10.1007/s10980-014-0082-3

Halley, J. M., Monokrousos, N., Mazaris, A. D., Newmark, W. D., and Vokou, D. (2016). Dynamics of extinction debt across five taxonomic groups. Nature Communications 7(1), 12283. https://doi.org/10.1038/ncomms12283

Hammer, Ø., Harper, D. A. T., and Ryan, P. D. (2001). PAST: paleontological statistics software package for education and data analysis. Palaeontologia Electronica 4, 1—9.

Hamrick, J. L., and Godt, M. J. W. (1996). Effects of life history traits on genetic diversity in plant species. Philosophical Transactions of the Royal Society of London. Series B, Biological Sciences 351, 1291–1298.

Helsen, K., Acharya, K. P., Brunet, J., Cousins, S. A. O., Decocq, G., Hermy, M., et al. (2017). Biotic and abiotic drivers of intraspecific trait variation within plant populations of three herbaceous plant species along a latitudinal gradient. BMC Ecology 17(1), 38. https://doi.org/10.1186/s12898-017-0151-y

Hoffmann, A. A., and Sgrò, C. M. (2011). Climate change and evolutionary adaptation. Nature 470(7335), 479–485. https://doi.org/10.1038/nature09670

IPCC (2007). Climate Change 2007: Synthesis Report. Contribution of Working Groups I, II and III to the Fourth Assessment Report. Intergovernmental Panel on Climate Change, Geneva, Switzerland.

Jenkins, D. G., Carey, M., Czerniewska, J., Fletcher, J., Hether, T., Jones, A., et al. (2010). A meta-analysis of isolation by distance: Relic or reference standard for landscape genetics? Ecography 33, 315–320. https://doi.org/10.1111/j.1600-0587.2010.06285.x

Kennard, W. C., Phillips, R. L., Porter, R. A., and Grombacher, A. W. (2000). A comparative map of wild rice (*Zizania palustris* L. 2n=2x=30). Theoretical and Applied Genetics 101(5–6), 677–684. https://doi.org/10.1007/s001220051530

Lande, R., and Shannon, S. (1996). The role of genetic variation in adaptation and population persistence in a changing environment. Evolution 50, 434–437.

Landguth, E. L., and Schwartz, M. K. (2014). Evaluating sample allocation and effort in detecting population differentiation for discrete and continuously distributed individuals. Conservation Genetics 15(4), 981–992. https://doi.org/10.1007/s10592-014-0593-0

Lee, C.-R., and Mitchell-Olds, T. (2011). Quantifying effects of environmental and geographical factors on patterns of genetic differentiation. Molecular Ecology 20(22), 4631–4642. https://doi.org/10.1111/j.1365-294X.2011.05310.x

Li, Y., Zhang, X.-X., Mao, R.-L., Yang, J., Miao, C.-Y., Li, Z., et al. (2017). Ten Years of Landscape Genomics: Challenges and Opportunities. Frontiers in Plant Science 8, 2136. https://doi.org/10.3389/fpls.2017.02136

Liu, B., Piao, H., Zhao, F., Zhao, J., and Zhao, R. (1999). Production and molecular characterization of rice lines with introgressed traits form a wild species *Zizania latifolia* (Griseb.). J. Genet. Breed. 53, 279–284.

Liu, J. G., Dong, Y., Xu, H., Wang, D. K., and Xu, J. K. (2007). Accumulation of Cd, Pb and Zn by 19 wetland plant species in constructed wetland. Journal of Hazardous Materials 147(3), 947–953. doi: 10.1016/j.jhazmat.2007.01.125

Lu, Y., Waller, D. M., and David, P. (2005). Genetic variability is correlated with population size and reproduction in American wild-rice (*Zizania palustris* var. Palustris, Poaceae) populations. American Journal of Botany 92(6), 990–997. https://doi.org/10.3732/ajb.92.6.990

Luque, S., Saura, S., and Fortin, M.-J. (2012). Landscape connectivity analysis for conservation: Insights from combining new methods with ecological and genetic data. Landscape Ecology 27(2), 153–157. https://doi.org/10.1007/s10980-011-9700-5

Manel, S., and Holderegger, R. (2013). Ten years of landscape genetics. Trends in Ecology & Evolution 28(10), 614–621. https://doi.org/10.1016/j.tree.2013.05.012

Manel, S., Gugerli, F., Thuiller, W., Alvarez, N., Legendre, P., Holderegger, R., Gielly, L., et al. (2012). Broad-scale adaptive genetic variation in alpine plants is driven by temperature and precipitation. Molecular Ecology 21(15), 3729–3738. https://doi.org/10.1111/j.1365-294X.2012.05656.x

Mantel, N. (1967). The detection of disease clustering and a generalized regression approach. Cancer Res. 27, 209–220.

Marrot, P., Garant, D., and Charmantier, A. (2017). Multiple extreme climatic events strengthen selection for earlier breeding in a wild passerine. Philosophical Transactions of the Royal Society B: Biological Sciences 372(1723), 20160372. https://doi.org/10.1098/rstb.2016.0372

McRae, B. H. (2006). Isolation by resistance. Evolution 60(8), 1551–1561. https://doi.org/10.1111/j.0014-3820.2006.tb00500.x

McRae, B. H., Dickson, B. G., Keitt, T. H., and Shah, V. B. (2008). Using circuit theory to model connectivity in ecology, evolution, and conservation. Ecology 89(10), 2712–2724. https://doi.org/10.1890/07-1861.1

Meirmans, P. G. (2012). The trouble with isolation by distance. Molecular Ecology 21(12), 2839–2846. https://doi.org/10.1111/j.1365-294X.2012.05578.x

Meirmans, P. G., and Van Tienderen, P. H. (2004). genotype and genodive: Two programs for the analysis of genetic diversity of asexual organisms. Molecular Ecology Notes 4(4), 792–794. https://doi.org/10.1111/j.1471-8286.2004.00770.x

Morente-López, J., García, C., Lara-Romero, C., García-Fernández, A., Draper, D., and Iriondo, J. M. (2018). Geography and Environment Shape Landscape Genetics of Mediterranean Alpine Species *Silene ciliata* Poiret. (Caryophyllaceae). Frontiers in Plant Science 9, 1698. https://doi.org/10.3389/fpls.2018.01698

Münzbergová, Z., Hadincová, V., Skálová, H., and Vandvik, V. (2017). Genetic differentiation and plasticity interact along temperature and precipitation gradients to determine plant performance under climate change. Journal of Ecology 105(5), 1358–1373. https://doi.org/10.1111/1365-2745.12762

Oksanen, J., Blanchet, F. G., Kindt, R., Legendre, P., Minchin, R. B., et al. (2018). vegan: Community Ecology Package. R package version 2.5-3. https://CRAN.R-project.org/package=vegan.

Orsini, L., Vanoverbeke, J., Swillen, I., Mergeay, J., and De Meester, L. (2013). Drivers of population genetic differentiation in the wild: Isolation by dispersal limitation, isolation by adaptation and isolation by colonization. Molecular Ecology 22(24), 5983–5999. https://doi.org/10.1111/mec.12561

Peakall, R., and Smouse, P. E. (2012). GenAlEx 6.5: Genetic analysis in Excel. Population genetic software for teaching and research--an update. Bioinformatics 28(19), 2537–2539. https://doi.org/10.1093/bioinformatics/bts460

Peng, S.-L., You, W.-H., Qi, G., and Yang, F.-L. (2013). Nitrogen and Phosphorus Uptake Capacity and Resource Use of Aquatic Vegetables Floating Bed in the Eutrophicated Lake. 2013 Third International Conference on Intelligent System Design and Engineering Applications 994–997. https://doi.org/10.1109/ISDEA.2012.237

Pritchard, J. K., Stephens, M., and Donnelly, P. (2000). Inference of population structure using multilocus genotype data. Genetics 155(2), 945–959.

Pykälä, J. (2019). Habitat loss and deterioration explain the disappearance of populations of threatened vascular plants, bryophytes and lichens in a hemiboreal landscape. Global Ecology and Conservation 18, e00610. https://doi.org/10.1016/j.gecco.2019.e00610

Qin, B. (2008). Lake Taihu, China: Dynamics and environmental change. Dordrecht: Springer.

Quan, Z., Pan, L., Ke, W., Liu, Y., and Ding, Y. (2009). Sixteen polymorphic microsatellite markers from *Zizania latifolia* Turcz. (Poaceae). Molecular Ecology Resources 9(3), 887–889. https://doi.org/10.1111/j.1755-0998.2008.02357.x

Richards, C. M., Antolin, M. F., Reilley, A., Poole, J., and Walters, C. (2007). Capturing genetic diversity of wild populations for ex situ conservation: Texas wild rice (*Zizania texana*) as a model. Genetic Resources and Crop Evolution 54(4), 837–848. https://doi.org/10.1007/s10722-006-9167-4

Rosenberg, N. A. (2003). distruct: A program for the graphical display of population structure. Molecular Ecology Notes 4(1), 137–138. https://doi.org/10.1046/j.1471-8286.2003.00566.x

Rousset, F. (1997). Genetic Differentiation and Estimation of Gene Flow from F-Statistics Under Isolation by Distance. Genetics 145(4), 1219.

Rousset, F., and Ferdy, J.-B. (2014). Testing environmental and genetic effects in the presence of spatial autocorrelation. Ecography 37(8), 781–790. https://doi.org/10.1111/ecog.00566

Santamaría, L. (2002). Why are most aquatic plants widely distributed? Dispersal, clonal growth and small-scale heterogeneity in a stressful environment. Acta Oecologica 23(3), 137–154. https://doi.org/10.1016/S1146-609X(02)01146-3

Schneider, S., Roessli, D., and Excoffier, L. (2000). Arlequin: A Software for Population Genetics Data Analysis, Version 2.0. Geneva, Switzerland: Genetics and Biometry Laboratory, Department of Anthropology, University of Geneva.

Segelbacher, G., Cushman, S. A., Epperson, B. K., Fortin, M.-J., Francois, O., Hardy, O. J., et al. (2010). Applications of landscape genetics in conservation biology: Concepts and challenges. Conservation Genetics 11(2), 375–385. https://doi.org/10.1007/s10592-009-0044-5

Sexton, J. P., Hangartner, S. B., and Hoffmann, A. A. (2014). Genetic isolation by environment or distance: which pattern of gene flow is most common?: special section. Evolution 68(1), 1–15. https://doi.org/10.1111/evo.12258

Shafer, A. B. A., and Wolf, J. B. W. (2013). Widespread evidence for incipient ecological speciation: A meta-analysis of isolation-by-ecology. Ecology Letters 16(7), 940–950. https://doi.org/10.1111/ele.12120

Shen, W., Song, C., Chen, J., Fu, Y., Wu, J., and Jiang, S. (2011). Transgenic Rice Plants Harboring Genomic DNA from *Zizania latifolia* Confer Bacterial Blight Resistance. In Rice Science 18(1), 17–22. https://doi.org/10.1016/S1672-6308(11)60003-6.

Slatkin, M. (1985). Gene flow in natural populations. Annu. Rev. Ecol. Syst. 16, 393–430.

Slatkin, M., and Barton, N. H. (1989). A comparison of three indirect methods for estimating average levels of gene flow. Evolution 43(7), 1349–1368. https://doi.org/10.1111/j.1558-5646.1989.tb02587.x

Sork, V. L., Nason, J., Campbell, D. R., and Fernandez, J. F. (1999). Landscape approaches to historical and contemporary gene flow in plants. Trends in Ecology & Evolution 14(6), 219–224. https://doi.org/10.1016/S0169-5347(98)01585-7

Su, H., Qu, L.-J., He, K., Zhang, Z., Wang, J., Chen, Z., and Gu, H. (2003). The Great Wall of China: A physical barrier to gene flow? Heredity 90(3), 212–219. https://doi.org/10.1038/sj.hdy.6800237

Šurinová, M., Hadincová, V., Vandvik, V., and Münzbergová, Z. (2019). Temperature and precipitation, but not geographic distance, explain genetic relatedness among populations in the perennial grass Festuca rubra. Journal of Plant Ecology 12(4), 730–741. https://doi.org/10.1093/jpe/rtz010

Tucker, J. M., Schwartz, M. K., Truex, R. L., Wisely, S. M., and Allendorf, F. W. (2014). Sampling affects the detection of genetic subdivision and conservation implications for fisher in the Sierra Nevada. Conservation Genetics 15(1), 123–136. https://doi.org/10.1007/s10592-013-0525-4

van Etten, J. (2017). R package gdistance: Distances and routes on geographical grids. Journal of Statistical Software 76(13), 1–21. doi: 10.18637/jss.v076.i13.

Van Oosterhout, C., Hutchinson, W. F., Wills, D. P. M., and Shipley, P. (2004). micro-checker: Software for identifying and correcting genotyping errors in microsatellite data. Molecular Ecology Notes 4(3), 535–538. https://doi.org/10.1111/j.1471-8286.2004.00684.x

van Strien, M. J., Holderegger, R., and Van Heck, H. J. (2015). Isolation-by-distance in landscapes: Considerations for landscape genetics. Heredity 114(1), 27–37. https://doi.org/10.1038/hdy.2014.62

Verity, R., and Nichols, R. A. (2016). Estimating the Number of Subpopulations (K) in Structured Populations. Genetics 203(4), 1827–1839. https://doi.org/10.1534/genetics.115.180992.

Wagutu, G. K., Njeri, H. K., Fan, X. R., and Chen, Y. Y. (2020). Development and transferability of SSR primers in the wild rice *Zizania latifolia* (Poaceae). Plant Science Journal 38(1), In Press.

Wang, H. M., Wu, G. L., Jiang, S. L., Huang, Q. N., Feng, B. H., Hunag, C. G., et al. (2015). Genetic diversity of *Zizania latifolia* Griseb. from Poyang lake basin based on SSR and ISSR analysis. Journal of Plant Genetic Resources 16(1), 133–141.

Wang, I. J., and Bradburd, G. S. (2014). Isolation by environment. Molecular Ecology 23(23), 5649–5662. https://doi.org/10.1111/mec.12938

Wang, M. X., Zhang, H. L., Zhang, D. L., Qi, Y. W., Fan, Z. L., Li, D. Y., et al. (2008). Genetic structure of *Oryza rufipogon* Griff. In China. Heredity 101(6), 527–535. https://doi.org/10.1038/hdy.2008.61

Wang, T., Wang, Z., Xia, F., and Su, Y. (2016). Local adaptation to temperature and precipitation in naturally fragmented populations of *Cephalotaxus oliveri*, an endangered conifer endemic to China. Scientific Reports 6(1), 25031.https://doi.org/10.1038/srep25031

Wang, Y., Huang, L., and Fan, L. (2013). Main agronomic traits, domestication and breeding of Gu (*Zizania latifolia*). Journal of Zhejiang University (Agriculture and Life Sciences) 39(6) 629–635. DOI: ISSN: 1008-9209 CN: 33-1247/S.

Wang, Z., Wu, J., Madden, M., and Mao, D. (2012). China’s Wetlands: Conservation Plans and Policy Impacts. AMBIO 41(7), 782–786. https://doi.org/10.1007/s13280-012-0280-7

Wright, S. (1943). Isolation by distance. Genetics 28 (2), 114–138.

Wright, S. (1969). Evolution and the genetics of populations: The theory of gene frequencies. The University of Chicago Press, Chicago, Illinois.

Wu. S., Yang, Q., and Zheng, D. (2003). Comparative study on eco-geographic regional systems between China and USA. Acta Geographica Sinica 58(5), 686–694.

Wu, Z., Xu, X., Zhang, J., Wiegleb, G., and Hou, H. (2019). Influence of environmental factors on the genetic variation of the aquatic macrophyte Ranunculus subrigidus on the Qinghai-Tibetan Plateau. BMC Evolutionary Biology 19(1), 228. https://doi.org/10.1186/s12862-019-1559-0

Xu, X. W., Wu, J. W., Qi, M. X., Lu, Q. X., Lee, P. F., Lutz, S., et al. (2015). Comparative phylogeography of the wild-rice genus *Zizania* (Poaceae) in eastern Asia and North America. Am J Bot. 102(2):239–47. doi: 10.3732/ajb.1400323.

Xu, X., Ke, W., Yu, X., Wen, J., and Ge, S. (2008). A preliminary study on population genetic structure and phylogeography of the wild and cultivated *Zizania latifolia* (Poaceae) based on Adh1a sequences. Theoretical and Applied Genetics 116(6), 835–843. https://doi.org/10.1007/s00122-008-0717-3

Xu, X., Walters, C., Antolin, M. F., Alexander, M. L., Lutz, S., Ge, S., et al. (2010). Phylogeny and biogeography of the eastern Asian–North American disjunct wild-rice genus (*Zizania* L., Poaceae). Molecular Phylogenetics & Evolution 55(3), 1008–1017. doi: 10.1016/j.ympev.2009.11.018.

Xu, Z., and Zhou, G. (2017). Identification and Control of Common Weeds. Dordrecht: Springer Netherlands.

Yu, C., Likun, L., Xiuyun, L., Wanli, G., and Bao, L. (2006). Isolation and characterization of a set of disease resistance-gene analogs (RGAs) from wild rice, *Zizania latifolia* Griseb. I. Introgression, copy number lability, sequence change, and DNA methylation alteration in several rice–Zizania introgression lines. Genome 49(2), 150–158. doi: 10.1139/G05-097.

Zhao, Y., Song, Z., Zhong, L., Li, Q., Chen, J., and Rong, J. (2019). Inferring the Origin of Cultivated *Zizania latifolia*, an Aquatic Vegetable of a Plant-Fungus Complex in the Yangtze River Basin. Frontiers in Plant Science 10, 1406.https://doi.org/10.3389/fpls.2019.01406

Zhao, Y., Vrieling, K., Liao, H., Xiao, M., Zhu, Y., Rong, J., et al. (2013). Are habitat fragmentation, local adaptation and isolation-by-distance driving population divergence in wild rice *Oryza rufipogon* ? Molecular Ecology 22(22), 5531–5547. https://doi.org/10.1111/mec.12517

Zhao, Y., Zhong, L., Zhou, K., Song, Z., Chen, J., and Rong, J. (2018). Seed characteristic variations and genetic structure of wild *Zizania latifolia* along a latitudinal gradient in China: Implications for neo-domestication as a grain crop. AoB PLANTS 10(6). https://doi.org/10.1093/aobpla/ply072

Zhou, S., Wang, C., Yang, H., Wang, G., Wang, Y., and Li, J. (2007). Growth of *Zizania latifolia* and *Acorus calamus* in sewage and their effect on sewage purification. Chin. J. App. Environ. Biol. 13, 454e457 (in Chinese with English abstract).

